# The molecular mechanism of Hsp90-induced oligomerization of Tau

**DOI:** 10.1101/614289

**Authors:** Sabrina Weickert, Magdalena Wawrzyniuk, Laura John, Stefan G. D. Rüdiger, Malte Drescher

## Abstract

Aggregation of the microtubule-associated protein Tau is a hallmark of Alzheimer’s disease with Tau oligomers suspected as the most toxic agent. Tau is a client of Hsp90, though it is unclear whether and how the chaperone massages the structure of intrinsically disordered Tau. Using electron paramagnetic resonance, we extract structural information from the very broad conformational ensemble of Tau: Tau in solution is highly dynamic and polymorphic, though ‘paper-clip’-shaped by long-range contacts. Interaction with Hsp90 promotes an open Tau conformation, which we identify as the molecular basis for the formation of small Tau oligomers by exposure of the aggregation-prone repeat domain to other Tau molecules. At the same time, formation of Tau fibrils is inhibited. We therefore provide the nanometer-scale zoom into chaperoning an amyloid client, highlighting formation of oligomers as the consequence of this biologically relevant interaction.

## Main Text

Tau is an intrinsically disordered protein (IDP) known to bind to and stabilize microtubules (MTs) and regulate axonal transport in its physiological function (*1-3*). In pathology, filamentous aggregates of Tau constitute a hallmark of neurodegenerative diseases, among them Alzheimer’s disease (AD) (*4*). Both MT-binding and self-aggregation of Tau are mediated by the Tau repeat domain (Tau-RD) consisting of 4 imperfect repeats in the longest Tau isoform (Fig. 1A) (*5, 6*).

**Figure 1.**
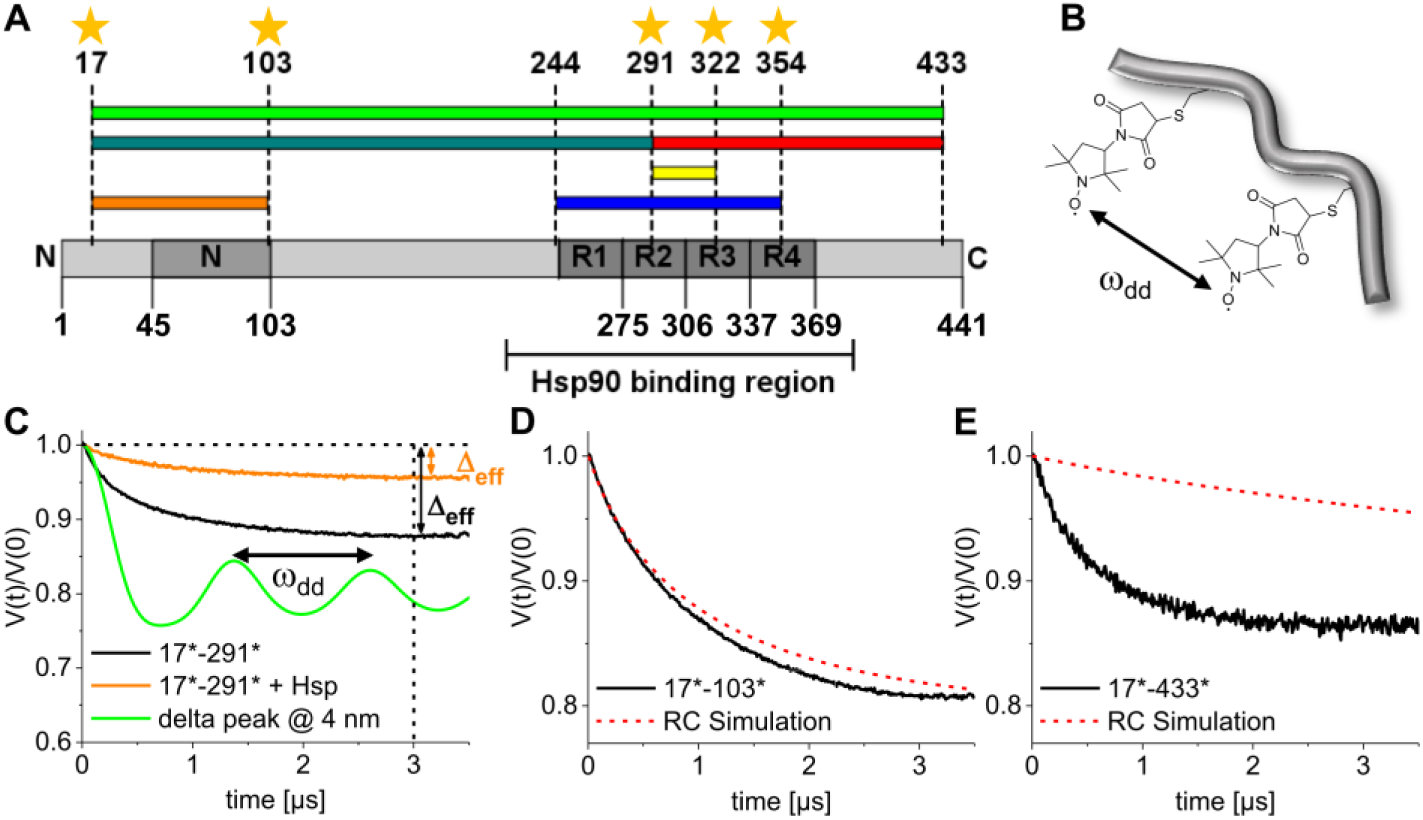
DEER with modulation depth-based data analysis allows to monitor the conformational ensemble of Tau. (A) Tau domain organization (R1-R4: pseudo-repeats, N: two N-terminal inserts). Stars indicate labeling positions in singly spin-labeled Tau derivatives. Colored bars depict sequences spanned by labels in doubly spin-labeled Tau. (B) 3-maleimido proxyl spin label side chains attached to cysteines. (C) Modulations with the dipolar modulation frequency ω_dd_ characterize a DEER time trace calculated for a delta peak at 4 nm (green). Experimental intramolecular DEER time traces recorded for Tau-17*-291* are modulation-free in the absence (black) and presence (orange) of Hsp90 indicating broad distributions of ω_dd_ and thus a broad conformational ensemble. Effective modulation depths Δ_eff_ at t = 3μs provide information about the Tau conformational ensemble without or with Hsp90. (D) A random coil model is in reasonable agreement with DEER results for several labeled stretches of Tau, e.g., Tau-17*-103*. (E) Experimental results, e.g., for Tau-17*-433* suggest a considerably larger vicinity of spin labels than the RC simulation predicts, consistent with a ‘paper-clip’ solution ensemble of Tau.

While Tau in solution is generally disordered and highly dynamic, long-range interactions mediate folding-back of both termini onto Tau-RD, resulting in an overall ‘paper-clip’ arrangement of monomeric Tau (*7, 8*). In filamentous aggregates of Tau, Tau-RD forms the ordered filamental core, while N- and C-terminal regions remain a disordered ‘fuzzy coat’ (*9, 10*). Tau filaments exist in different morphologies with striking differences in the fold of the filamental cores, which are probably disease-specific (*11, 12*). Although fibrils have long been considered the neurotoxic species, neuronal death appears rather to be caused by prefibrillar soluble aggregates and oligomers of Tau (*13, 14*), which are also considered responsible for spreading Tau pathogenicity from cell-to-cell in a prion-like concept (*15, 16*).

The molecular chaperone Hsp90 (*17, 18*) initiates proteasomal degradation (*19-21*) and induces oligomerization of Tau (*22-24*). Tau-RD is part of the Hsp90/Tau interaction interface (*25*). While insight into the molecular mechanism of the emergence of toxic Tau oligomers is highly relevant in the context of neuropathology, its structural principle is elusive.

The lack of a defined three-dimensional fold of IDPs like Tau makes their structural characterization challenging. Electron paramagnetic resonance (EPR) spectroscopy in combination with site-directed spin labeling (SDSL) has proven powerful in the investigation of IDPs and their aggregation behavior also in the presence of diverse interaction partners (*26-31*). EPR spectroscopy (i) provides information on the side-chain dynamics of a single residue (*32*). Dipolar spectroscopy, i.e., DEER (double electron-electron resonance) spectroscopy (ii) gives access to distance information in the nm range between two spin labels by measuring their magnetic dipolar interaction frequency ω_dd_ (*33-35*). Here, we exploit the combination of these approaches to investigate the molecular mechanism of Hsp90-induced Tau oligomerization.

We genetically engineered Tau derivatives containing one or two cysteines at specific sites and performed thiol-specific spin labeling (Fig. 1B). A range of biochemical and biophysical assays showed that these spin-labeled Tau alterations maintained the structural and functional integrity of the wild type protein (Fig. S1-S4, Table S1).

Next, we set out to characterize the structural properties of Tau by obtaining long-range intramolecular distance information with DEER on doubly labeled Tau. Typical experimental DEER form factors for Tau are shown in Fig. 1C (full data Fig. S5) in comparison to simulated data for a hypothetical, well-defined distance. In contrast to the latter, the experimental traces for Tau showed no distinct modulations, indicating a broad distribution of spin-spin distances and thus implying a vast conformational ensemble of Tau in solution.

For these experimental DEER traces, the standard method for DEER data analysis fails and the extraction of precise distance distributions is precluded (*36, 37*). We compared the experimental DEER traces with simulated traces calculated from a random coil (RC) model (Fig. S6). For certain spin-labeled stretches of Tau, e.g., Tau-17*-103*, the RC model agreed well with the experimental results (Fig. 1D, S6), indicating an RC-like structural ensemble in these Tau segments. However, the RC model cannot describe the whole DEER data set: For Tau-17*-291* and Tau-17*-433*, the deviation between experiment and RC model indicates a considerable contribution from Tau conformations more compact than RC (Fig. 1E).

Hinderberger *et al*. proposed a data analysis procedure (*36, 37*), which we adapted for analyzing the broad conformational ensemble of a large IDP like Tau. We evaluated the DEER data using the effective modulation depth Δ_eff_, which is the signal decay of the DEER time trace at time t = 3 μs (Fig. 1C). While a DEER trace in the absence of Hsp90 delivers a reference Δ_eff_ value for each Tau sample, the change ΔΔ_eff_ upon addition of Hsp90 characterizes transitions in the conformational equilibrium: Negative ΔΔ_effs_ indicate an increase in spin-spin separation, while positive Δ_effs_ are consistent with the spins coming into closer proximity of each other (Fig. S7). This allows extracting distance information from DEER traces not analyzable in the conventional way.

The systematic analysis of the experimental Δ_eff_ values supports the ‘paper-clip’ model proposed on the basis of FRET and NMR experiments for Tau in solution, where N- and C-termini are in proximity to each other and Tau-RD in an overall more compact fold than RC (*7, 8*). On the one hand these results demonstrate the capacity of the Δ_eff_ approach for obtaining structural information from DEER traces reflecting vast protein ensembles, while on the other hand they define the ‘paper-clip’ as a reference structural ensemble of Tau in solution.

It has been shown that Hsp90 induces oligomerization of Tau *fragments* (*22*). Here, we analyzed the oligomerization behavior of full-length Tau by density gradient centrifugation (Fig. 2). Pure Tau was mainly found in its monomeric form, while heparin induced the formation of mature fibrils. In the presence of Hsp90 the amount of *small* oligomeric Tau species increased. Strikingly, the formation of high-molecular weight Tau aggregates and fibrils was prevented in the presence of Hsp90. Electron micrographs of K18 Tau fragments in the presence of Hsp90 also show the formation of small protein conglomerates, while fibril formation is prevented.(*38*)

**Figure 2.**
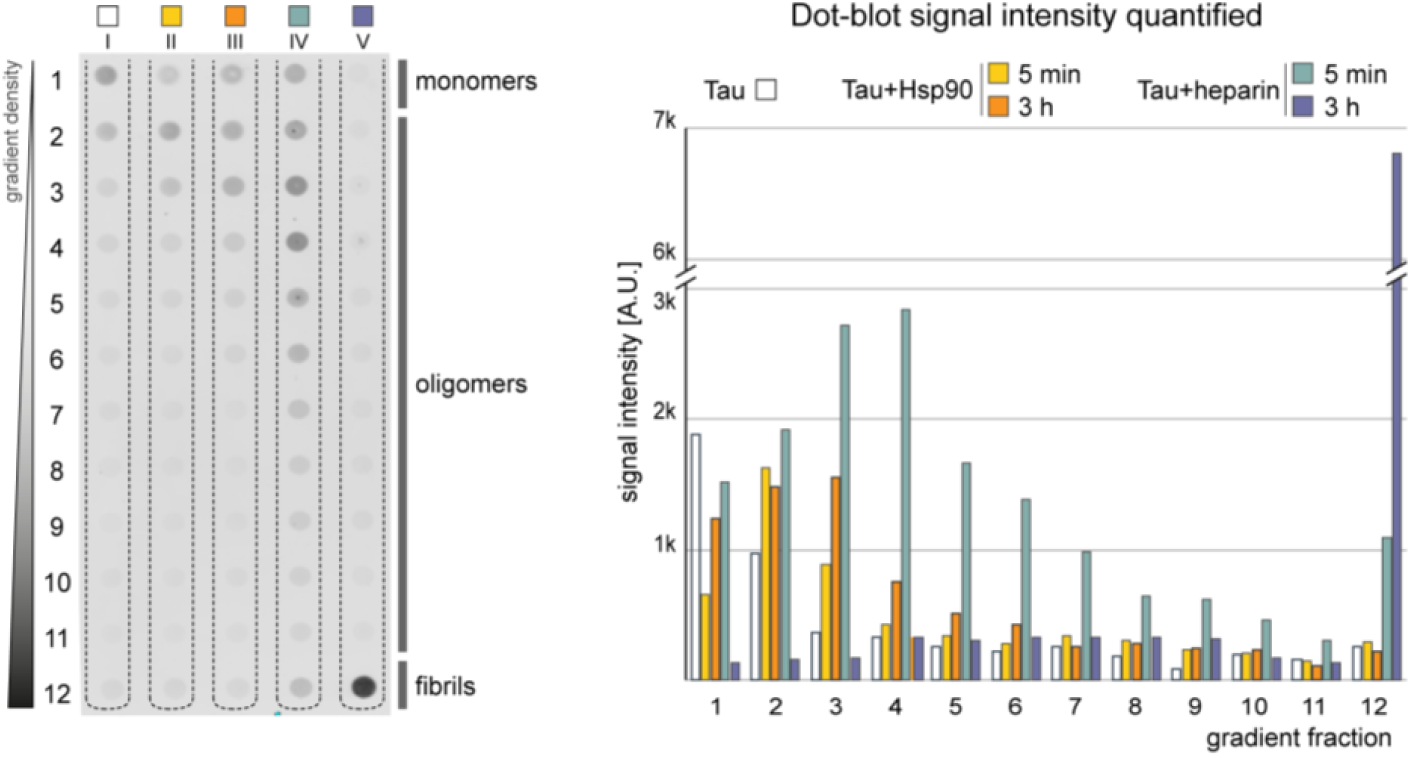
Hsp90 promotes the formation of small oligomeric species of Tau. Dot blot summarizing the results of density gradient centrifugation and quantification: Pure Tau is mostly monomeric. Heparin induces formation of high-molecular weight fibrils. Hsp90 leads to an increase in small Tau oligomeric species, while formation of fibrils is prohibited.

To identify the oligomerization-domain in Tau relevant for Hsp90-induced oligomerization, we performed intermolecular DEER measurements using singly spin-labeled Tau: Upon oligomerization Δ_eff_ would increase locally where inter-Tau contacts are established. We observed very small Δ_eff_ values for all Tau derivatives in the absence of Hsp90 (Fig. 3) indicating only minor subpopulations of oligomeric Tau species. Addition of Hsp90 lead to a considerable increase in Δ_eff_ for Tau-322* and Tau-354*, depicted as difference values ΔΔ_eff_. This suggests that the oligomerization interface is located in Tau-RD, and specifically in R3/R4. Strikingly, Tau oligomerization initiates in the same Tau region responsible for AD fibril formation as well as Hsp90 binding (*11, 25*). This is remarkable, as it suggests that the same stretch of Tau mediating fibril formation (*25*) is addressed by Hsp90 to promote the formation of oligomers.

**Figure 3.**
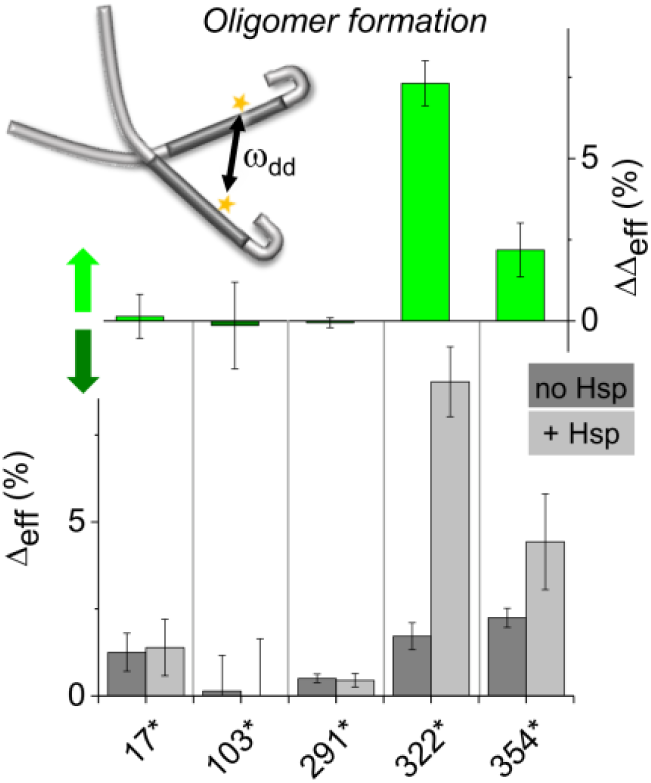
Oligomerization is mediated by R3/R4. Information about intermolecular Tau/Tau interactions obtained with DEER of singly spin-labeled Tau in the absence (dark gray) and presence (light gray) of Hsp90. Non-zero Δ_eff_s represent small amounts of non-monomeric Tau in the absence of Hsp90. Hsp90 increased Δ_eff_s for Tau-322* and Tau-354* (light green bars) in accordance with an increase in Tau oligomers mediated by R3/R4 of Tau-RD.

The dynamic properties of Tau in solution and with Hsp90 are reported by EPR spectra of spin-labeled Tau side chains. Generally, we observed rather fast rotational dynamics with rotational correlation times τ_corr_ around 1 ns (Fig. 4A). This is in accordance with Tau presiding in a largely unstructured state with a broad conformational ensemble and a high degree of dynamical disorder (*26*). Addition of Hsp90 induced only subtle changes in the spectra (Fig. S8), indicating that dynamic disorder in Tau persists also when bound. The generally still fast dynamics in the Tau spectra demonstrate the transient nature of the Tau/Hsp90 complex. We determined the half lifetime of the Tau/Hsp90 complex by quartz crystal microbalance (QCM) affinity measurements at ∼10 s, which is typical for transient protein-protein interactions (Fig. S9) (*39*). The Tau/Hsp90 complex appears to be characterized by transient interactions between individual residues, involving a structural multiplicity of Tau.

**Figure 4.**
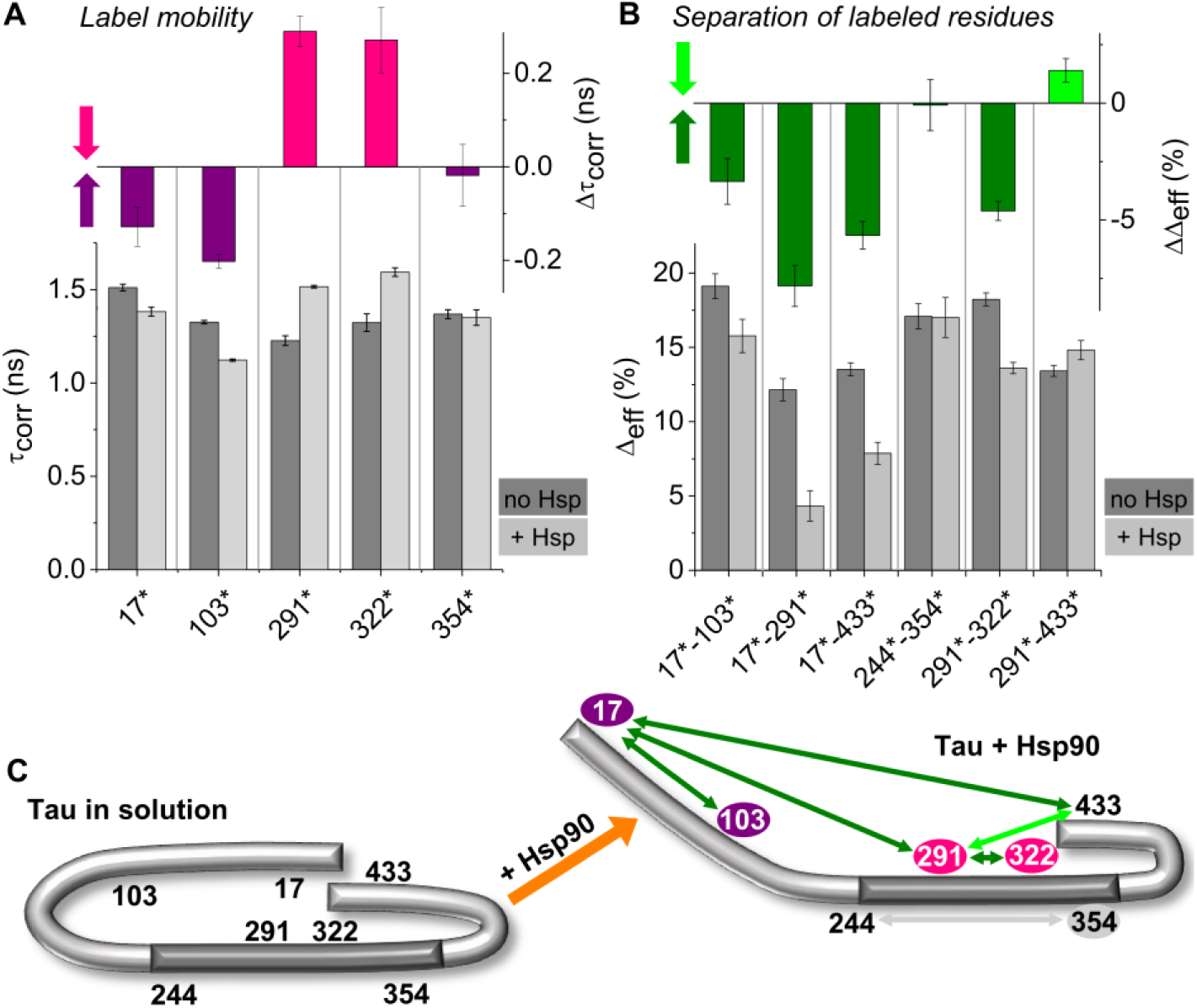
The Tau conformational ensemble opens up upon interaction with Hsp90. (A) Local side-chain dynamics accessed by EPR of singly spin-labeled Tau derivatives: Rotational correlation times τ_corr_ determined in the absence (dark gray) and presence (light gray) of Hsp90 and respective changes Δτ_corr_ (purple/pink) are shown. Arrows indicate a decrease (pink) or increase (purple) in side-chain mobility at the respective site. (B) Intramolecular distance information obtained by DEER spectroscopy with doubly spin-labeled Tau derivatives: Effective modulation depths Δ_eff_ determined in the absence (dark gray) and presence (light gray) of Hsp90 and respective changes ΔΔ_eff_ (light/dark green) are shown. Arrows indicate a decrease (light green) or increase (dark green) in spin label separation. (C) Structural ensemble model of Tau in the absence (left) and presence of Hsp90 (right) derived from EPR data in (A) and (B). In the absence of Hsp90, Tau adopts a ‘paper-clip’ shape with both termini folded back onto Tau-RD. In the presence of Hsp90 the N-terminus folds outwards, thereby uncovering Tau-RD.

We observed local restrictions of the reorientational mobility for spin-labeled side chains Tau-291* and Tau-322* in the presence of Hsp90. Both residues are located in Tau-RD, which has been identified as the Hsp90-binding region before (*25*). Thus, the altered dynamics are attributed to direct Tau/Hsp90 interaction, while also oligomer formation might restrict side chain dynamics of Tau-322*.

Spin label mobilities increased for Tau-17* and Tau-103* upon addition of Hsp90, indicating that these side chains gain a larger conformational space. Thus, one might speculate that the N-terminus detaches from Tau-RD upon binding of Hsp90, opening up the ‘paper-clip’ fold.

To elucidate the structural influence of Hsp90 on the Tau conformational ensemble, we performed DEER spectroscopy of doubly spin-labeled Tau. DEER traces remained modulation-free upon addition of Hsp90. Thus, dynamic disorder prevails in Tau also when interacting with the chaperone (Fig. 1C). Addition of Hsp90 changed Δ_eff_ values, indicating a shift in the conformational equilibrium of Tau: A pronounced increase in the average spin-spin separation occurred for Tau-17*-291* and Tau-17*-433*. This indicates that the N-terminus detaches from both Tau-RD and the C-terminus and folds outwards, opening up the ‘paper-clip’ (Fig. 4C). ΔΔ_eff_ values suggested a slight stretching of N-terminal Tau between Tau-17*-103* and of Tau-RD in the region between Tau-291*-322* in R2/R3, while the overall dimension of Tau-RD between Tau-244*-354* remained unchanged. While individual repeat sequences, e.g., R2/R3 expanded while accommodating Hsp90, there seems to be considerable flexibility in the remaining Tau-RD for preserving its overall dimension. A similar structural reorganization of Tau towards an open conformation was reported upon binding to tubulin, where stretches between individual repeats expanded, while the overall dimension of Tau-RD was unaffected (*40*). Our results suggest a molecular mechanism for Tau oligomerization: Binding to Hsp90 opens the compact Tau solution structure, exposing Tau-RD residues and presenting them to other Tau molecules. As the Tau/Hsp90 complex is of a transient nature, oligomerization of Tau molecules may then occur via exposed Tau-RD.

With a combination of magnetic resonance experiments and biochemical assays, we gained detailed insight into the conformational ensemble of the IDP Tau in the presence of the molecular chaperone Hsp90. While DEER spectroscopy is routinely used to extract distance restraints in well-ordered proteins or small segments of disordered proteins, we demonstrated in the current study, how DEER in combination with a sophisticated data analysis approach can be profitably employed in obtaining structural information from a highly polymorphic and dynamic conformational ensemble of a large, full-length IDP. This approach enabled us to shed light on the molecular mechanism of the Tau/Hsp90 interaction.

Probing Tau conformation in solution revealed a ‘paper-clip’ arrangement of the domains (Fig. 4). The N-terminal domain forms long-range interactions with the aggregation-prone domain, in line with previous findings (*22-24*). The flexible N-terminus may, therefore, preclude aberrant Tau/Tau interactions, resulting in the high solubility of the full-length protein. Indeed, Tau fragments lacking the N-terminal region, and therefore exposing the aggregation prone repeat region, feature drastically accelerated aggregation rates, such as Tau-RD (*41*). Remarkably, Tau-RD is less soluble than the full-length protein despite having a higher net charge (pI 9.67 for Tau-RD *vs*. 8.24 for full-length) (*41, 42*). Hsp90 binds to Tau-RD, consistent with previous findings by us (*25*). The Tau/Hsp90 complex is distinguished by transient protein-protein interactions with Tau remaining conformationally heterogeneous and dynamically disordered, which is characteristic for ‘fuzzy’ complexes (*43, 44*) and was also described for a ternary Hsp90/PPIase/Tau complex (*45*). In the light of the large extension of the Tau/Hsp90 interaction interface (*25*) it is consistent that single residues contribute only little to the overall binding between Hsp90 and Tau, enabling a high rate of unbinding (*43, 44*). This is a typical way for Hsp90 to bind its client proteins (*46, 47*). Also Tau typically retains a high degree of conformational dynamics, at least for protein segments, when in complex with various interaction partners including heparin, tubulin, MTs and in Tau fibrils (*9, 26, 40, 48, 49*). Binding to Hsp90 induces a conformational opening of ‘paper-clip’-Tau (7, 8), leading to exposure of Tau-RD.

Hsp90-binding does not alter the global dimension of Tau-RD, similar as Tau bound to tubulin (*40*). Heparin, in contrast, compacts Tau-RD (*50, 51*). This suggests that Hsp90 stabilizes Tau in a conformation different from the heparin-induced aggregation-prone one, but similar to a conformation which might enable productive binding to MTs (*19*). It remains to be seen whether the Hsp90-bound shape may be similar to the fibril structure in AD. The Tau/Hsp90 interaction promotes self-aggregation after opening the compact Tau solution structure, probably due to exposure of the aggregation-prone Tau-RD. Remarkably, Hsp90-seeded oligomers exhibit intermolecular DEER signal solely for the variants sampling the two C-terminal repeats R3 and R4 (Fig. 3) that mediate the early steps of Tau aggregation (*26*) and eventually constitute the core of the paired helical filament of AD (*6, 11*). However, the continuation to fibril formation is blocked in the presence of Hsp90, which may foster the generation of Tau oligomers. The conformational transition and opening up of Tau is therefore the structural basis for subsequent formation of deleterious oligomers in the presence of Hsp90.

## Acknowledgments

We thank Stefan Stoll for extensive discussion concerning DEER data analysis. We thank Elke Deuerling for fruitful discussions. We thank Madelon Maurice for collaboration in the Initial Training Network “WntsApp” (No. 608180), supported by Marie-Curie Actions of the 7th Framework programme of the EU. We thank Dr. Davide Proverbio and Dr. Teodor Aastrup for the guidance in the QCM measurements at Attana AB. We appreciate the help of Luca Ferrari with density gradient analysis. This work was supported by the Deutsche Forschungsgemeinschaft (SFB 969; project C03). S.G.D.R. was further supported by the Internationale Stichting Alzheimer Onderzoek (ISAO; project “Chaperoning Tau Aggregation”; No. 14542) and a ZonMW TOP grant (“Chaperoning Axonal Transport in neurodegenerative disease”; No. 91215084).

## References

1. E. M. Mandelkow, E. Mandelkow, Biochemistry and cell biology of tau protein in neurofibrillary degeneration. Cold Spring Harb Perspect Med 2, a006247 (2012).

2. Y. Wang, E. Mandelkow, Tau in physiology and pathology. Nat. Rev. Neurosci. 17, 5–21 (2016).

3. E. H. Kellogg et al., Near-atomic model of microtubule-tau interactions. Science 360, 1242–1246 (2018).

4. M. Goedert, Alzheimer’s and Parkinson’s diseases: The prion concept in relation to assembled Abeta, tau, and alpha-synuclein. Science 349, 1255555 (2015).

5. T. Guo, W. Noble, D. P. Hanger, Roles of tau protein in health and disease. Acta Neuropathol. 133, 665–704 (2017).

6. T. Crowther, M. Goedert, C. M. Wischik, The repeat region of microtubule-associated protein tau forms part of the core of the paired helical filament of Alzheimer’s disease. Ann. Med. 21, 127–132 (1989).

7. S. Jeganathan, M. von Bergen, H. Brutlach, H.-J. Steinhoff, E. Mandelkow, Global Hairpin Folding of Tau in Solution. Biochemistry 45, 2283–2293 (2006).

8. M. D. Mukrasch et al., Structural polymorphism of 441-residue tau at single residue resolution. PLoS Biol. 7, e34 (2009).

9. C. M. Wischik et al., Structural characterization of the core of the paired helical filament of Alzheimer disease. Proc. Natl. Acad. Sci. U.S.A. 85, 4884–4888 (1988).

10. S. Wegmann, I. D. Medalsy, E. Mandelkow, D. J. Müller, The fuzzy coat of pathological human Tau fibrils is a two-layered polyelectrolyte brush. Proc. Natl. Acad. Sci. USA 110, E313–E321 (2013).

11. A. W. P. Fitzpatrick et al., Cryo-EM structures of tau filaments from Alzheimer’s disease. Nature 547, 185–190 (2017).

12. B. Falcon et al., Structures of filaments from Pick’s disease reveal a novel tau protein fold. Nature 561, 137–140 (2018).

13. C. A. Lasagna-Reeves et al., Identification of oligomers at early stages of tau aggregation in Alzheimer’s disease. FASEB J. 26, 1946–1959 (2012).

14. C. A. Lasagna-Reeves et al., Tau oligomers impair memory and induce synaptic and mitochondrial dysfunction in wild-type mice. Mol. Neurodegener. 6, 39 (2011).

15. F. Clavaguera et al., Transmission and spreading of tauopathy in transgenic mouse brain. Nat. Cell Biol. 11, 909–913 (2009).

16. L. Liu et al., Trans-Synaptic Spread of Tau Pathology In Vivo. PLoS One 7, e31302 (2012).

17. F. U. Hartl, A. Bracher, M. Hayer-Hartl, Molecular chaperones in protein folding and proteostasis. Nature 475, 324 (2011).

18. F. H. Schopf, M. M. Biebl, J. Buchner, The HSP90 chaperone machinery. Nat. Rev. Mol. Cell Biol. 18, 345–360 (2017).

19. F. Dou et al., Chaperones increase association of tau protein with microtubules. Proc. Natl. Acad. Sci. U.S.A. 100, 721–726 (2003).

20. C. A. Dickey et al., The high-affinity HSP90-CHIP complex recognizes and selectively degrades phosphorylated tau client proteins. J. Clin. Invest. 117, 648–658 (2007).

21. A. D. Thompson et al., Analysis of the Tau-Associated Proteome Reveals That Exchange of Hsp70 for Hsp90 Is Involved in Tau Degradation. ACS Chem. Biol. 7, 1677–1686 (2012).

22. E. Tortosa et al., Binding of Hsp90 to tau promotes a conformational change and aggregation of tau protein. J. Alzheimer’s Dis. 17, 319–325 (2009).

23. L. J. Blair et al., Accelerated neurodegeneration through chaperone-mediated oligomerization of tau. J. Clin. Invest. 123, 4158–4169 (2013).

24. L. B. Shelton et al., Hsp90 activator Aha1 drives production of pathological tau aggregates. Proc. Natl. Acad. Sci. U.S.A. 114, 9707–9712 (2017).

25. G. E. Karagöz et al., Hsp90-Tau Complex Reveals Molecular Basis for Specificity in Chaperone Action. Cell 156, 963–974 (2014).

26. A. Pavlova et al., Protein structural and surface water rearrangement constitute major events in the earliest aggregation stages of tau. Proc. Natl. Acad. Sci. USA 113, E127–E136 (2016).

27. F.-X. Theillet et al., Structural disorder of monomeric α-synuclein persists in mammalian cells. Nature 530, 45–50 (2016).

28. M. Robotta, J. Cattani, J. C. Martins, V. Subramaniam, M. Drescher, Alpha-Synuclein Disease Mutations Are Structurally Defective and Locally Affect Membrane Binding. J. Am. Chem. Soc. 139, 4254–4257 (2017).

29. Y. Fichou, M. Vigers, A. K. Goring, N. A. Eschmann, S. Han, Heparin-induced tau filaments are structurally heterogeneous and differ from Alzheimer’s disease filaments. Chem. Commun. 54, 4573–4576 (2018).

30. N. Le Breton et al., Exploring intrinsically disordered proteins using site-directed spin labeling electron paramagnetic resonance spectroscopy. Front. Mol. Biosci. 2, 21 (2015).

31. S. Weickert, J. Cattani, M. Drescher, in Electron Paramagnetic Resonance: Volume 26. (The Royal Society of Chemistry, 2019), vol. 26, pp. 1–37.

32. W. L. Hubbell, D. S. Cafiso, C. Altenbach, Identifying conformational changes with site-directed spin labeling. Nat. Struct. Biol. 7, 735–739 (2000).

33. A. Milov, A. Ponomarev, Y. D. Tsvetkov, Electron-electron double resonance in electron spin echo: Model biradical systems and the sensitized photolysis of decalin. Chem. Phys. Lett. 110, 67–72 (1984).

34. M. Pannier, S. Veit, A. Godt, G. Jeschke, H. W. Spiess, Dead-time free measurement of dipole–dipole interactions between electron spins. J. Magn. Reson. 213, 316–325 (2000).

35. G. Jeschke, DEER distance measurements on proteins. Annu. Rev. Phys. Chem. 63, 419–446 (2012).

36. D. Kurzbach et al., Cooperative Unfolding of Compact Conformations of the Intrinsically Disordered Protein Osteopontin. Biochemistry 52, 5167–5175 (2013).

37. D. Kurzbach et al., Compensatory Adaptations of Structural Dynamics in an Intrinsically Disordered Protein Complex. Angew. Chem. Int. Ed. 53, 3840–3843 (2014).

38. L. Ferrari et al., Fibril formation rewires interactome of the Alzheimer protein Tau by π-stacking. bioRxiv, 522284 (2019).

39. J. R. Perkins, I. Diboun, B. H. Dessailly, J. G. Lees, C. Orengo, Transient protein-protein interactions: structural, functional, and network properties. Structure 18, 1233–1243 (2010).

40. A. M. Melo, J. Coraor, S. Elbaum-Garfinkle, A. Nath, E. Rhoades, A functional role for intrinsic disorder in the tau-tubulin complex. Proc. Natl. Acad. Sci. U.S.A. 113, 14336–14341 (2016).

41. S. Barghorn, E. Mandelkow, Toward a Unified Scheme for the Aggregation of Tau into Alzheimer Paired Helical Filaments. Biochemistry 41, 14885–14896 (2002).

42. N. D. Keul et al., The entropic force generated by intrinsically disordered segments tunes protein function. Nature 563, 584–588 (2018).

43. R. Sharma, Z. Raduly, M. Miskei, M. Fuxreiter, Fuzzy complexes: Specific binding without complete folding. FEBS Lett. 589, 2533–2542 (2015).

44. M. Fuxreiter, Fold or not to fold upon binding - does it really matter? Curr. Opin. Struct. Biol. 54, 19–25 (2019).

45. J. Oroz et al., Structure and pro-toxic mechanism of the human Hsp90/PPIase/Tau complex. Nat. Commun. 9, 4532 (2018).

46. M. Radli, S. G. D. Rüdiger, Dancing with the Diva: Hsp90-Client Interactions. J. Mol. Biol. 430, 3029–3040 (2018).

47. M. Taipale, D. F. Jarosz, S. Lindquist, HSP90 at the hub of protein homeostasis: emerging mechanistic insights. Nat. Rev. Mol. Cell Biol. 11, 515–528 (2010).

48. H. Kadavath et al., Tau stabilizes microtubules by binding at the interface between tubulin heterodimers. Proc. Natl. Acad. Sci. U.S.A. 112, 7501–7506 (2015).

49. M. Martinho et al., Two Tau binding sites on tubulin revealed by thiol-disulfide exchanges. Sci. Rep. 8, 13846 (2018).

50. S. Elbaum-Garfinkle, E. Rhoades, Identification of an aggregation-prone structure of tau. J. Am. Chem. Soc. 134, 16607–16613 (2012).

51. D. Fischer et al., Conformational changes specific for pseudophosphorylation at serine 262 selectively impair binding of tau to microtubules. Biochemistry 48, 10047–10055 (2009).

