## Supplementary Material for "The molecular mechanism of Hsp90-induced oligomerization of Tau"

### Abstract:

Aggregation of the microtubule-associated protein Tau is a hallmark of Alzheimer's disease with Tau oligomers suspected as the most toxic agent. Tau is a client of Hsp90, though it is unclear whether and how the chaperone massages the structure of intrinsically disordered Tau. Using electron paramagnetic resonance, we extract structural information from the very broad conformational ensemble of Tau: Tau in solution is highly dynamic and polymorphic, though 'paper-clip'-shaped by long-range contacts. Interaction with Hsp90 promotes an open Tau conformation, which we identify as the molecular basis for the formation of small Tau oligomers by exposure of the aggregation-prone repeat domain to other Tau molecules. At the same time, formation of Tau fibrils is inhibited. We therefore provide the nanometer-scale zoom into chaperoning an amyloid client, highlighting formation of oligomers as the consequence of this biologically relevant interaction.

### Materials and Methods

Protein expression and spin labeling

Hsp90 and Tau were prepared as described before (*1, 2*). Briefly, N-terminally FLAG-tagged (DYKDDDDK) human Tau (isoform F) was expressed in E. coli BL21* cells overnight at 24 °C. To facilitate purification, an additional cleavable 6xHis-Smt tag (MGHHHHHHGSDSEVNQEAKPEVKPEVKPETHINLKVSDGSSEIFFKIKKTTPLRRLM EAFAKRQGKEMDSLRFLYDGIRIQADQTPEDLDMEDNDIIEAHREQIGG) was added upstream of the FLAG-Tau sequence. Cells were then disrupted using microfluidizer EmulsiFlex-C5 (Avestin) in the lysis buffer (25 mM HEPES, pH 7.2, 150 mM NaCl, 5 mM β-mercaptoethanol, 1 cOmplete Protease Inhibitor tablet (Roche) per pellet) and afterwards heated for 20 min at 353 K before clearing the lysate by centrifugation and filtering through a 0.22 μm syringe filter (VWR, polypropylene). Tau was then purified on an ÄKTA purifier chromatography system (GE Healthcare) by affinity purification (POROS 20MC, Thermo Fisher Scientific; 50 mM HEPES, pH 7.2, 300 mM NaCl, 10 mM β-mercaptoethanol; elution by a 0-100% gradient up to 1 M imidazole (Sigma Aldrich) over 5 CV) followed by anion exchange chromatography (POROS 20HQ; 50 mM HEPES pH 7.5, 10 mM β-mercaptoethanol; elution by a 0-100% gradient up to 2 M KCl over 5 CV). The fractions containing the Tau fusion protein were pooled and treated for 12 h at 277 K with Ulp1 protease to remove the 6xHis-SUMO tag. Cleavage products were separated on ÄKTA by cation exchange chromatography (column POROS 20HS, buffers as for 20HQ column). Last step of purification was to ensure homogeneity of the protein by size exclusion chromatography (column HiLoad 26/60 Superdex 200 pg, GE Healthcare) in the destination buffer (25 mM HEPES, pH 7.5, 75 mM NaCl, 75 mM KCl, 5 mM DTT). Eluted fractions were concentrated by centrifugation using a Vivaspin column (molecular weight cutoff 10 kDa) in sample buffer (25 mM HEPES pH 7.5, 75 mM NaCl, 75 mM KCl, 1 mM TCEP, cOmplete Protease Inhibitor (Roche)), aliquoted and stored at 193 K. Protein purity was confirmed by SDS-PAGE. For spin labeling, Tau derivatives containing 1 or 2 cysteine residues were thawed on ice and centrifuged at 277 K (10 min, 16000*g) in order to isolate potential precipitates. Sample volumes and concentrations were adjusted to 500 μl and 100 μM with sample buffer using spin columns (molecular weight cutoff 10 kDa). Per cysteine a six-fold molar excess of 3-maleimido-proxyl spin label was added to the samples and incubated overnight in the dark at 277 K. After the labeling reaction, free spin label was removed in multiple washing steps with sample buffer and the resulting volume of the samples was adjusted to 200 μl. Protein concentrations were quantified using a BCA protein assay kit with a reference (Thermo Scientific). Spin concentrations were quantified using the reference-free spin counting module of the EMXnano (Bruker Biospin). Near quantitative spin labeling was confirmed under the applied conditions. Success and specificity of the labeling reaction were confirmed by FTMS+ESI-MS measurements on a LTQ Orbitrap Discovery spectrometer (Thermo Scientific). The integrity of spin-labeled Tau derivatives was confirmed by SDS-PAGE.

Circular-Dichroism (CD) measurements

CD measurements (190 to 260 nm) were carried out with a Jasco J815 spectropolarimeter (Jasco Instruments). Spectra were recorded at 293 K in demountable cuvettes with a path length of 0.5 mm (Hellma). The scanning speed was 100 nm/min with a bandwidth of 1.0 nm and a response time of 1 s. In each measurement, 6 scans were accumulated and averaged.

Raw spectra were subjected to baseline correction, subtraction of the buffer background spectrum and smoothing by the means-movement method. For secondary structure interpretation, experimental CD spectra were fitted with a linear combination of standard CD spectra (*3*). The fit parameters were systematically varied in order to determine the error margins within which acceptable fits were obtained.

Samples of 28 μM protein concentration were prepared with spin-labeled and non-spin-labeled Tau. Samples were measured in sample buffer and/or PBS buffer after buffer exchange (140 mM NaCl, 10 mM $Na_2HPO_4$/1.8 mM $KH_2PO_4$, pH 7.4, 2.7 mM KCl), as PBS allows detection at lower wavelenghts. The spectrum recorded with PBS buffer and all simulations were normalized to their minimum. The spectra recorded in sample buffer were normalized to the value of the simulation at the lowest evaluable wavelength (high tension (HT) = 550 V).

EPR sample preparation

Samples for continuous wave (cw) EPR and DEER measurements were prepared with deuterated sample buffer and had final Tau concentrations of 28 μM. Hsp90 was added where applicable at a concentration of 56 μM. For measurements at cryogenic temperatures, 20 % (vol/vol) [$d_8$]glycerol was added to the samples.

cw-EPR experiments

All cw EPR spectra were recorded with singly spin-labeled Tau derivatives at 293 K using a MiniScope MS5000 spectrometer (magnettech) equipped with a variable temperature unit TC-H04 (magnettech). Samples with a typical sample volume of 10 μl were loaded into glass capillaries (inner diameter 1.03 mm). Spectra were recorded using an incident microwave power of 3.16 mW, with a modulation amplitude of 400 mG and a sweep width of 200 G. Typically, the signal-to-noise ratio was improved by accumulating 90 scans with 60 s sweep time each.

The spectra were analyzed using homewritten Matlab scripts. Rotational correlation times $\tau_{corr}$ were obtained using Kivelson’s equation

$$\tau_{\mathrm{corr}} = 6{,}6 \cdot 10^{-10} \cdot \mathrm{W}_0 \left[ \sqrt{\frac{\mathrm{h}_0}{\mathrm{h}_{-1}}} + \sqrt{\frac{\mathrm{h}_0}{\mathrm{h}_{+1}}} - 2 \right]$$

where $W_0$ is the peak-to-peak line width of the central peak and $h_{-1}$, $h_0$ and $h_{+1}$ are the intensities of the high, center and low field peaks (*4*). The standard deviations of the rotational correlation times were determined by the Gaussian error propagation law assuming that the peak intensities are statistically independent (*5*).

DEER experiments

DEER measurements were performed at 50 K on doubly and singly spin-labeled Tau derivatives with a sample volume of 12.5 μl. Each sample was collected in a quartz capillary (inner diameter 1 mm) and shock-frozen in liquid nitrogen within 10 min incubation time on ice and stored at 193 K until measurement without unfreezing.

Pulsed Q-band (34 GHz) DEER experiments were performed using an ELEXSYS E580 spectrometer equipped with an EN 5107D2 Q-band resonator (both Bruker BioSpin) and a 10W solid state amplifier (HBH Microwave). The temperature was adjusted with a helium gas flow system (CF935, Oxford Instruments). The pulse sequence for the four-pulse, dead-time-free DEER experiments was: $\pi/2_{obs} - \tau_1 - \pi_{obs} - t' - \pi_{pump} - (\tau_1 + \tau_2 - t') - \pi_{obs} - \tau_2 -$ echo (*6*).

The resonator was overcoupled (Q ≈ 100) and pulse lengths were optimized for each experiment. Typical observer pulse lengths $\pi/2_{obs}$ and $\pi_{obs}$ were 22 to 26 ns and 44 to 52 ns, respectively. Typical pump pulse lengths $\pi_{pump}$ were 24 to 28 ns. The time separation $\tau_1$ was always 400 ns. The echo amplitude was recorded as a function of the dipolar evolution time t'. The pump frequency $\nu_{pump}$ was set to the center of the resonator dip, where the global maximum of the nitroxide EPR spectrum was positioned. The observer frequency $\nu_{obs}$ was shifted by 50 MHz towards lower frequencies to a local maximum of the nitroxide EPR spectrum. The shot repetition time was 4080 μs and total measurement time for each sample was typically 15-24 h.

DEER raw data were analyzed globally using the DD (Version 7B) software package for Matlab (*7, 8*). DEER traces were normalized to 1 at t = 0 and to the excitation bandwidth of a 28 ns pump pulse. All DEER traces were evaluated with a dipolar evolution time of 3.5 μs. Extraction of distance distributions was not possible because of the non-modulated nature of the experimental DEER traces. Instead, effective modulation depths $\Delta_{eff}$ were determined as a measure for the effective spin-spin separation as established by Hindeberger and coworkers at a dipolar evolution time $t = 3\mu s$ (*9, 10*). DEER traces were described by a fit $F(t) = O(t) \times E(t)$, where $O(t)$ is the DEER signal for all spins within a given nanoobject based on a Gaussian spin-spin distribution, and $E(t)$ is an exponential function describing the background component. The echo intensity V(t) at $t = 3\mu s$ was determined from the fit after removal of the background component. The effective modulation depth was determined as $\Delta_{eff} = \left(1 - V(t = 3\mu s)\right) * 100$. Error bars give the 2σ confidence level of the parameter uncertainties as determined by the DD software.

To ensure the reliability of data evaluation, we also used the DeerAnalysis2016 software package for Matlab for data analysis (*11*), where background correction of the raw data is performed before evaluation of the resulting form factor with model-free Tikhonov regularization. We used different background starts in the analysis, e.g., set manually to 2.5 μs or as determined by the DeerAnalysis2016 software. The results obtained for $\Delta_{eff}$ are very robust and rather independent of the data evaluation method.

Random coil model calculations

Tau is intrinsically disordered, which suggests random coil (RC) as a suitable model to describe the expected spatial separation between two spin-labeled sites in the Tau protein. In order to compare RC simulations with the experimental DEER data, we calculated DEER traces from simulated distance distributions P(r) as derived from an RC model. The characteristic distributions P(r) of the label-to-label distances were calculated according to an ideal chain model as

$$P(r) = 4\pi r^2 \left(\frac{3}{2\pi\langle r^2\rangle}\right)^{3/2} e^{-\frac{3r^2}{2\langle r^2\rangle}}.$$

The radius of gyration is linked to the average label-to-label distance $\sqrt{\langle r^2\rangle}$ by

$$R_g = \sqrt{\langle r^2\rangle}\Big/\sqrt{6},$$

and, according to Kohn and coworkers, can also be described for peptides as

$$R_g = 1.927\ \text{Å} * N^{0.6},$$

where N is the number of residues between the two spin-labeling positions including the labeled residues (*12*). From P(r), DEER traces were calculated using DeerAnalysis2016 with an inversion efficiency of 20% and compared to the DEER traces recorded experimentally (see Fig. 1, S7).

<u>Sucrose gradient ultracentrifugation and dot blot</u>
Tau samples incubated with Hsp90 or heparin (incubation times as indicated in Fig. 2, 37 °C, 160 rpm) were resolved by ultracentrifugation on sucrose density gradients. To prepare the gradients, Beckman ultracentrifugation 12 ml tubes were filled with 6 ml 40% sucrose solution and carefully topped with 6 ml 10% sucrose (sucrose, 25 mM HEPES pH 7.5, 75 mM NaCl, 75 mM KCl, filtered through 0.22 μm), and incubated for 3 h in horizontal position (tilted by 90°). Density gradients were subjected to dot blot analysis. After centrifugation, each sucrose gradient was divided into 12 equal fractions of increasing density. 250 μl of each fraction were applied onto a nitrocellulose membrane (0.1 μm, Sigma Aldrich) using a dot blot chamber (BioRad), followed by immunodetection and fluorimetric visualisation. After sample application, the membrane was removed from the chamber and blocked for 1 h with PBS blocking buffer (Odyssey). Tau content in each fraction was detected using antibody against FLAG-tag (M2, Sigma; 1 h at 1:10,000 dilution) in PBS (137 mM NaCl, 2.7 mM KCl, 8 mM $Na_2HPO_4$ and 2 mM $KH_2PO_4$, pH 7.4) with 50% of Odyssey blocking buffer. After washing (5 x 5 min with PBS), the membrane was incubated for 45 min with the Alexa-680 conjugated secondary antibody at the dilution of 1:20 000 (donkey anti-mouse, Invitrogen). The antibody was removed and the membrane was washed 5 x 5 min before detection. Prepared membrane was scanned on LiCor Odyssey CLx instrument with the 700 nm laser line. Blots were analyzed with ImageStudioLite LiCor software with the empty well selected for the background subtraction.

<u>Quartz crystal microbalance</u>
The low-non-specific binding (LNB) quartz crystal chip was coated with recombinant Hsp90 by amine coupling. Briefly, two chips were activated for 300 s with freshly prepared solution of 0.2 M 1-ethyl-3-(3-dimethylaminopropyl)-carbodiimide with 0.05 M sulfo-*N*-hydroxysuccinimide at the flow of 10 μl/min. 50 μM Hsp90 in sodium acetate buffer, pH 4.5 was injected on one of the chips for 300 s. Identical buffer missing the protein was used for the control chip. Coupling was terminated by 300 s injection of 1 M ethanoloamine, pH 8.5. Binding experiments were performed on Attana A200 C-Fast system (Attana AB) in PBS buffer with 5 mM DTT. Tau was injected in duplicate in seven concentrations ranging from 1 to 80 μg/ml on Hsp90-coated chip and empty crystal for the reference (Channel A and Channel B). The chip surface was regenerated with 10 mM glycine, pH 3.5, between every injection. The binding affinity and dynamics were calculated for the binding curve obtained by subtracting the values measured on an empty chip from values measured on an Hsp90-coated chip. A 1:2 binding model established Bmax1 of 87% and Bmax2 of 13%. The Tau-Hsp90 binding affinity and dynamics are the parameters for the major binding component, Bmax1.

**Figures S1 – S9**

**Figure S1.**

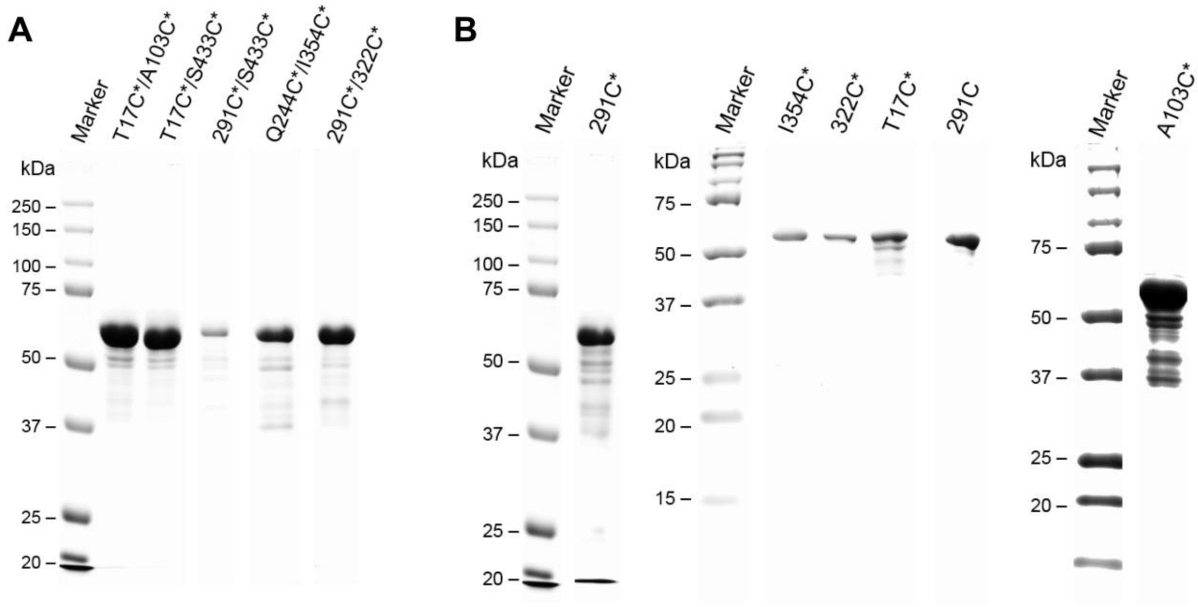

**SDS gels of spin-labeled Tau.** SDS-PAGE of doubly (**A**) and singly spin-labeled (**B**) Tau derivatives after completion of the overnight spin labeling reaction and the following washing procedure as used for EPR experiments. All samples are of reasonable purity and only gels that were overloaded with protein show a minor fraction of degradation products.

**Figure S2.**

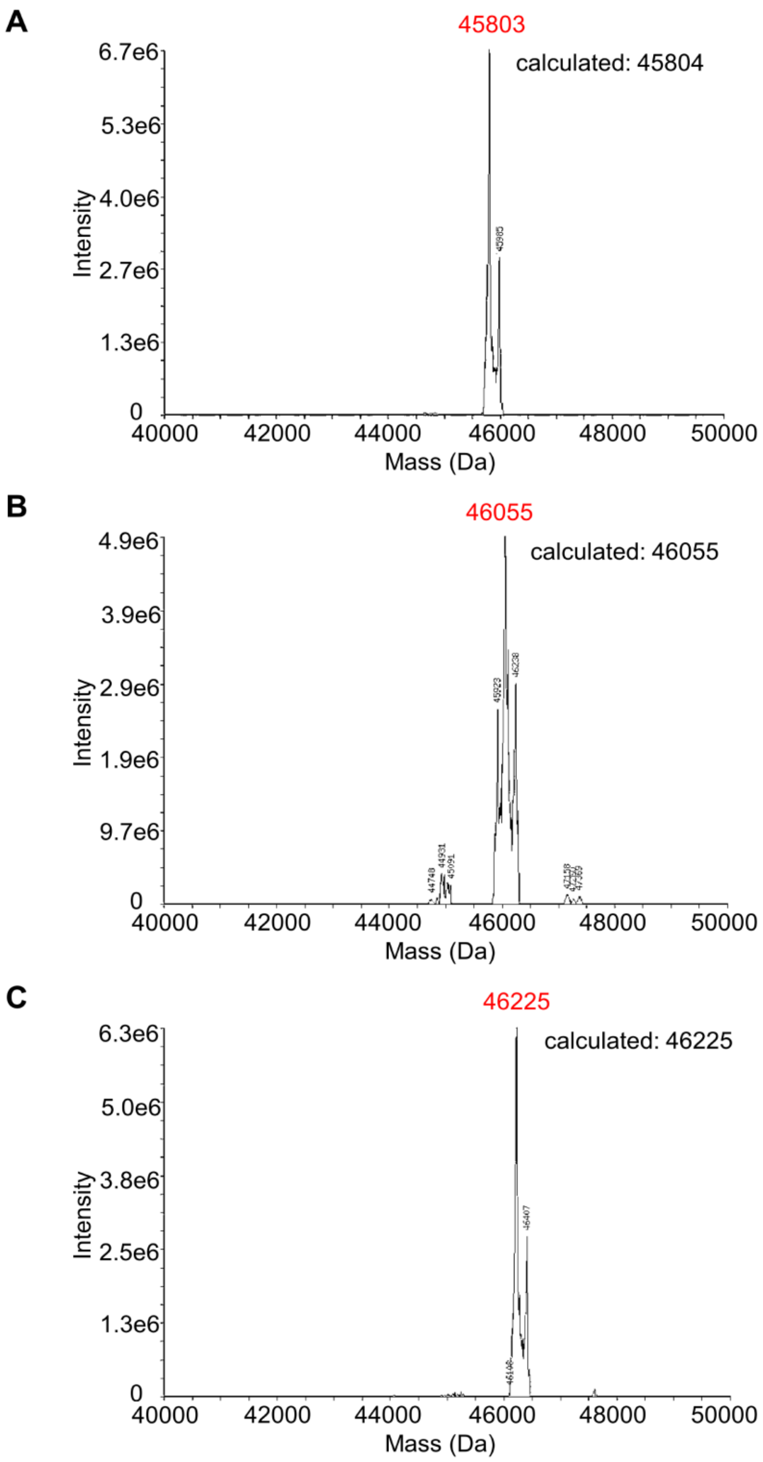

**Mass spectrometry of spin-labeled Tau.** FTMS+ESI-MS of unlabeled Tau-C17-C433 (**A**), singly spin-labeled Tau-291* (**B**) and doubly spin-labeled Tau-244*-354* (**C**). In all cases the main mass detected in the spectra coincides with the calculated mass of the respective Tau construct. No unlabeleld Tau is detected in the the spectrum of singly spin-labeled Tau and no unlabeled or singly spin-labeled Tau is present in the spectrum of doubly spin-labeled Tau.

**Figure S3.**

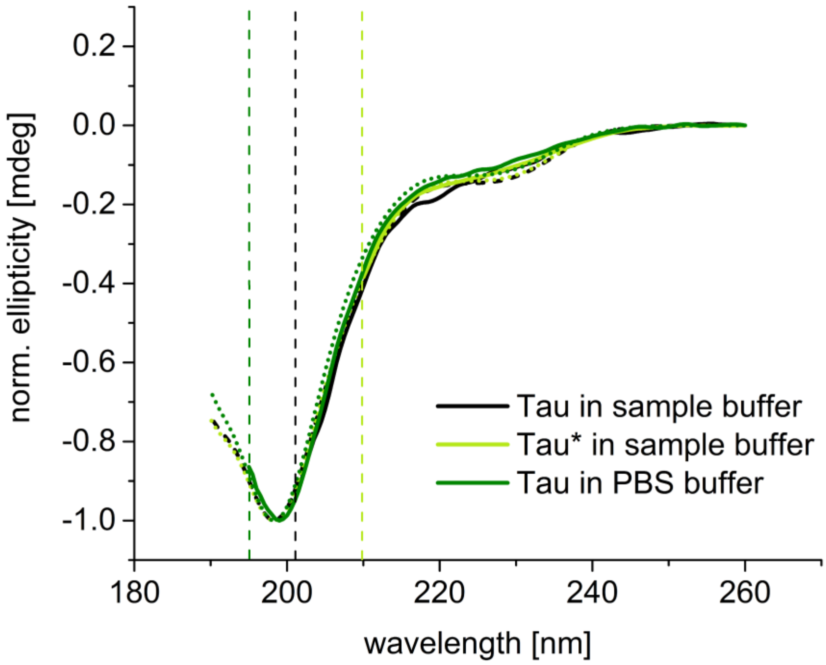

**CD spectra of Tau.** CD spectra of spin-labeled and non-spin-labeled Tau in sample buffer and PBS. All CD spectra were recorded with Tau-291$^{(*)}$-433$^{(*)}$. Irrespective of the degree of spin labeling or the choice of buffer all recorded CD spectra are basically identical. Corresponding simulations of the spectra are shown in Table S1. Simulations were obtained as a linear combination of standard CD spectra from distinct secondary structure elements (*3*). All experimental spectra can be described by simulations with very similar or identical parameters (see Table S1). The predominant contribution is from a random coil CD spectrum in all cases, which is in accordance with the intrinsically disordered character of Tau. Minor contributions from β-strand, β-turn and α-helix secondary structure enable a very good description of the experimental spectra by the simulation.

**Figure S4.**

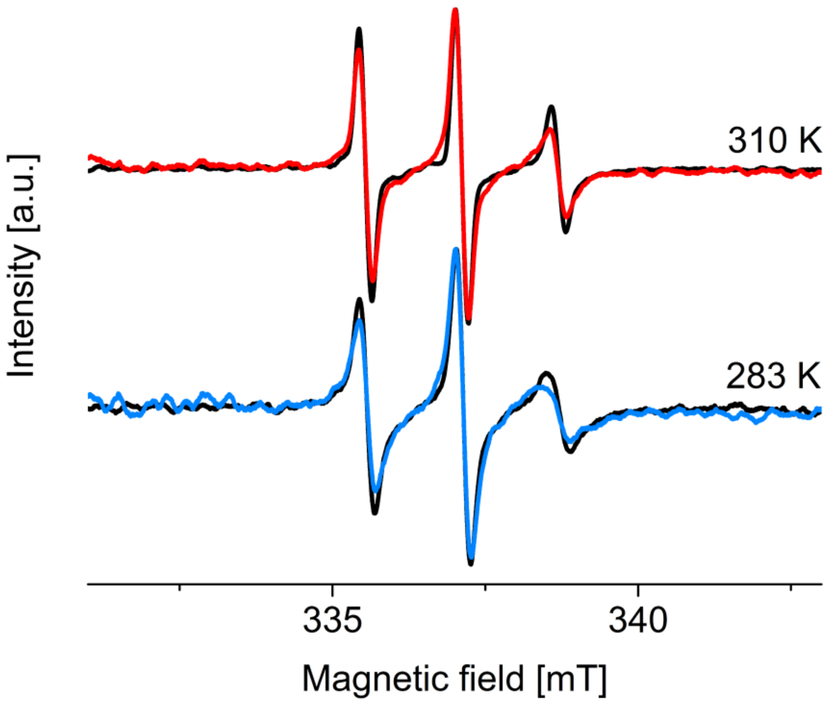

**Microtubule polymerization in the presence of spin-labeled Tau.** Cw EPR spectra of Tau-322* in buffer (black) or in the presence of tubulin recorded at 283 K and 310 K (blue and red, respectively). The spin label of the Tau derivative is located in the MTBR. In the presence of tubulin the spectral shapes report a decrease in spin label mobility as expected for Tau-322* interacting with tubulin. We conclude from these data that Tau constructs remain functional also after spin labeling with 3-maleimido-proxyl.

**Figure S5.**

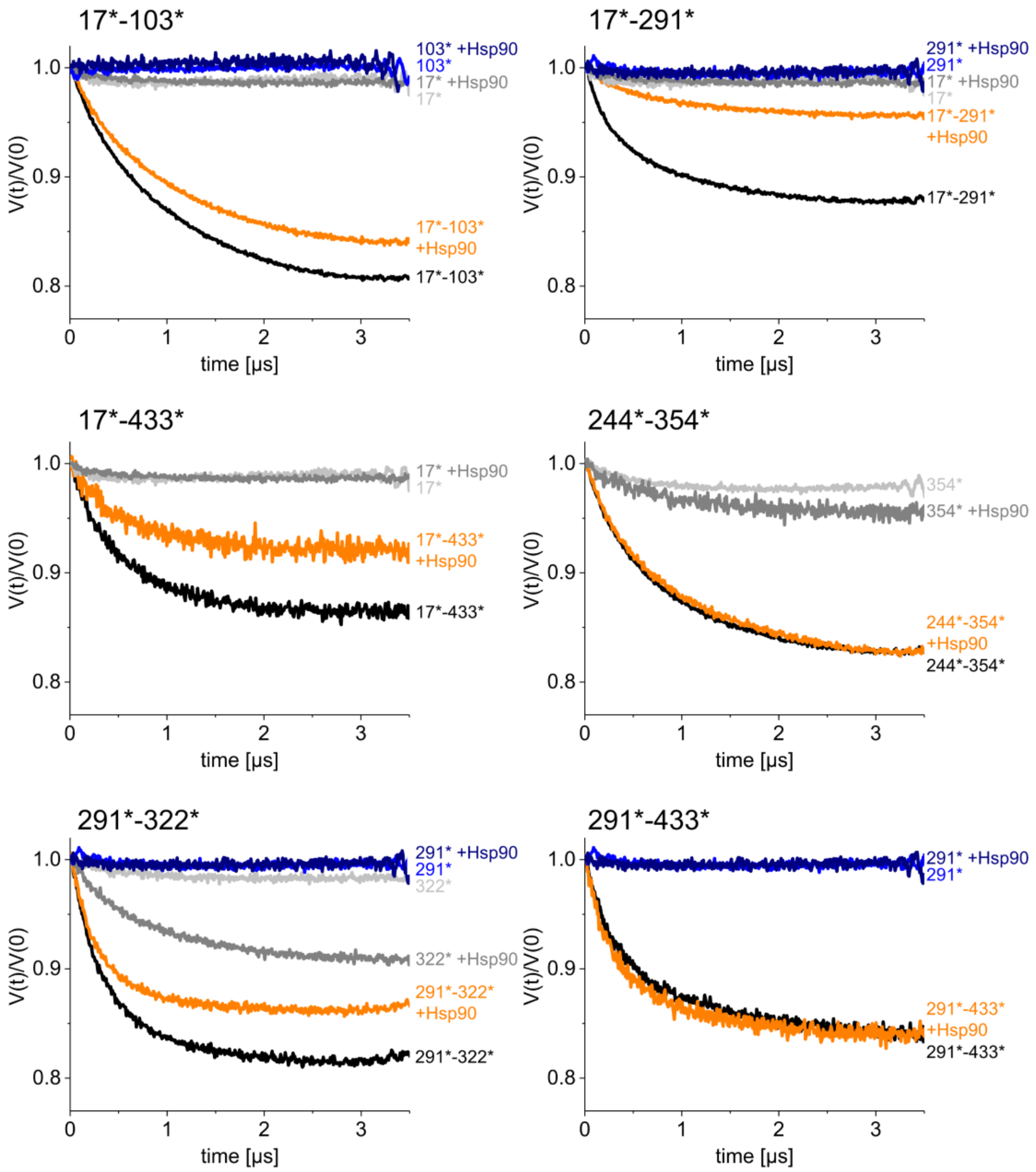

**DEER data.** Background-corrected DEER traces of doubly spin-labeled Tau in the absence (black) and presence of Hsp90 (orange) and corresponding DEER traces of singly spin-labeled Tau in the absence (light gray, blue) and presence (dark gray, dark blue) of Hsp90. All DEER traces are basically free of characteristic modulations with the dipolar interaction frequency $\omega_{dd}$, which is inversely proportional to $r^{-3}$, where r is the spin-spin distance. This indicates a very broad distribution of distances, which contribute to the experimental DEER curves. This is in accordance with a vast conformational ensemble of Tau in the absence and presence of Hsp90.

Classical extraction of a distance distribution from these modulation-free DEER traces fails. However, the parameter $\Delta_{eff}$ extracted at t = 3 μs allows to qualitatively assess the average spin-spin distance and analyze structural transitions in the Tau conformational ensemble upon addition of Hsp90.

**Figure S6.**

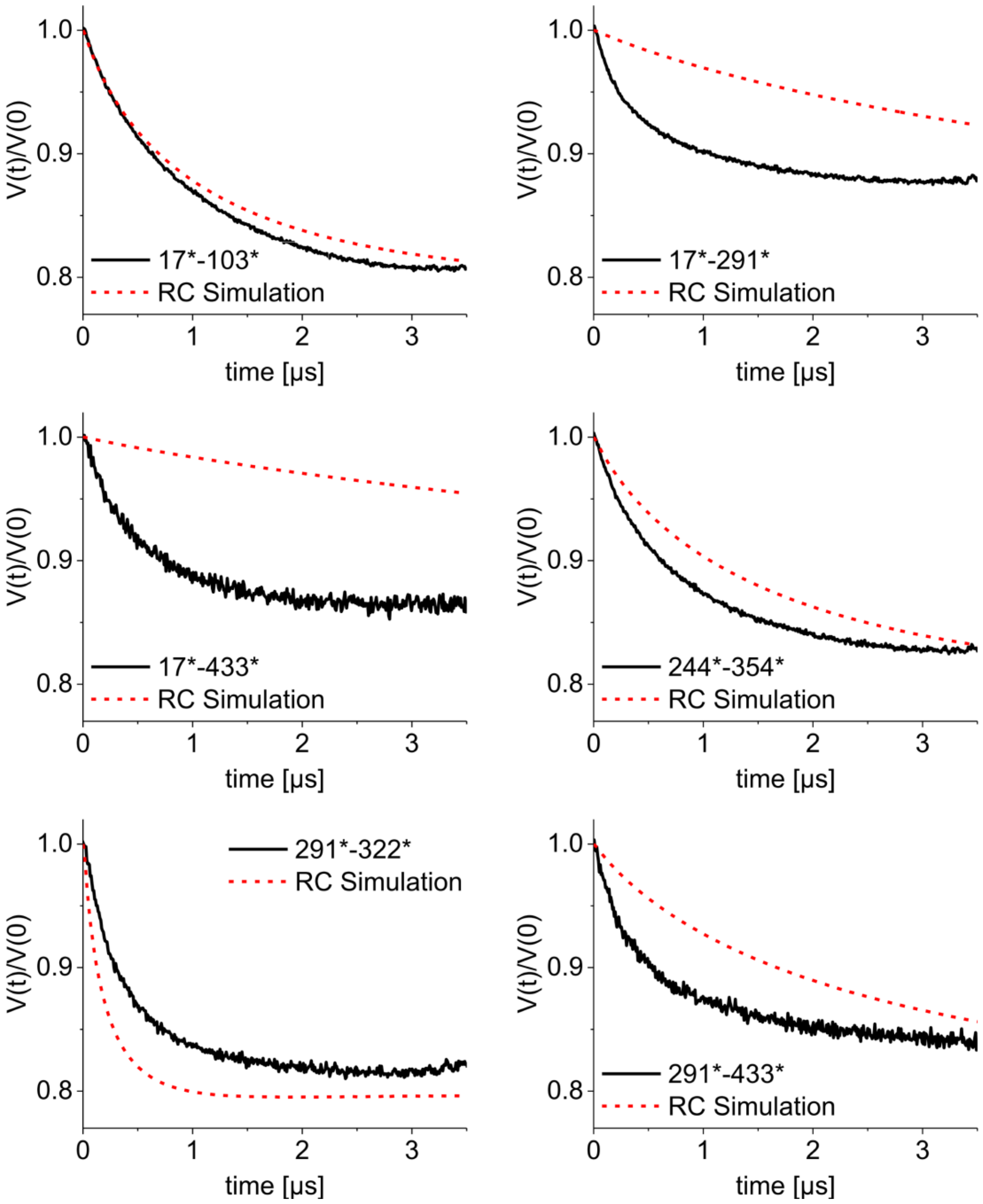

**Random coil model.** Comparison of DEER traces measured for doubly spin-labeled Tau derivatives (black) with DEER traces calculated from an RC model with the mean label-to-label distance derived from the number of residues located in between spin labels including the labeled residues (red dotted). While the dimensions of the N-terminal part (17*-103*) and the overall MTBR (244*-354*) match well with an RC model, it is obvious, that soluble Tau is more compact than RC between the N-terminus and the MTBR (17*-291*) and the N-terminus and the C-terminus (17*-433*). Also the conformation measured between the MTBR and the C-terminus (291*-433*) is much more compact than expected for an RC protein. On the other hand, R2 and R3 seem to be slightly more extended than predicted by an RC model, which is in accordance with the results of Melo et al. (*13*)

**Figure S7.**

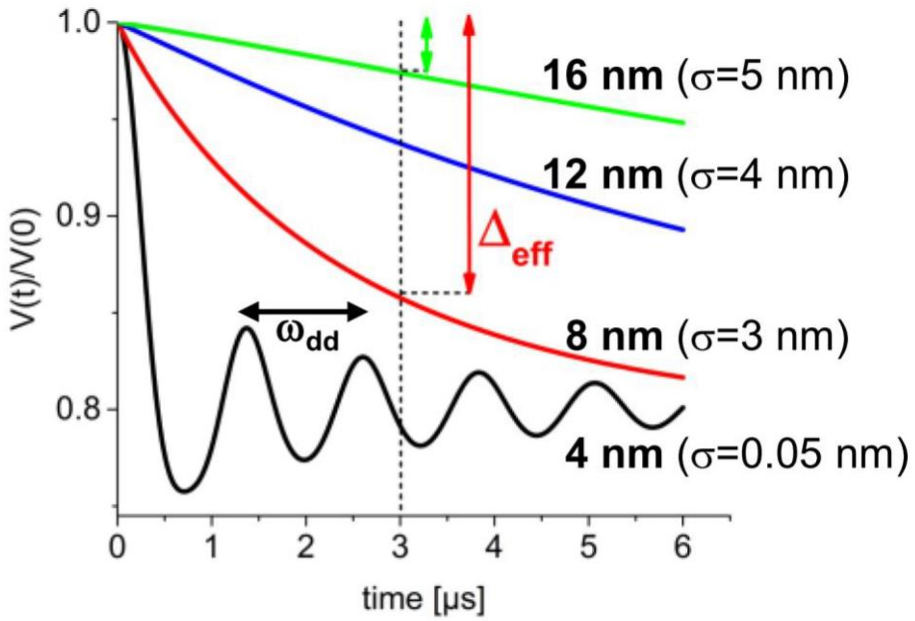

**$\Delta_{eff}$ as a measure for spin-spin separation.** Simulated DEER time traces calculated from Gaussian distance distributions as indicated. While a narrow 4 nm distance distribution (black) produces a DEER trace modulated with the dipolar interaction frequency $\omega_{dd}$, broad distance distributions (red, blue, green) are modulation-free. The effective modulation depth $\Delta_{eff}$ at t = 3 µs decreases with increasing average spin-spin distance $\langle r^2\rangle^{1/2}$ and can in this way be employed as a measure for the average interspin separation.

**Figure S8.**

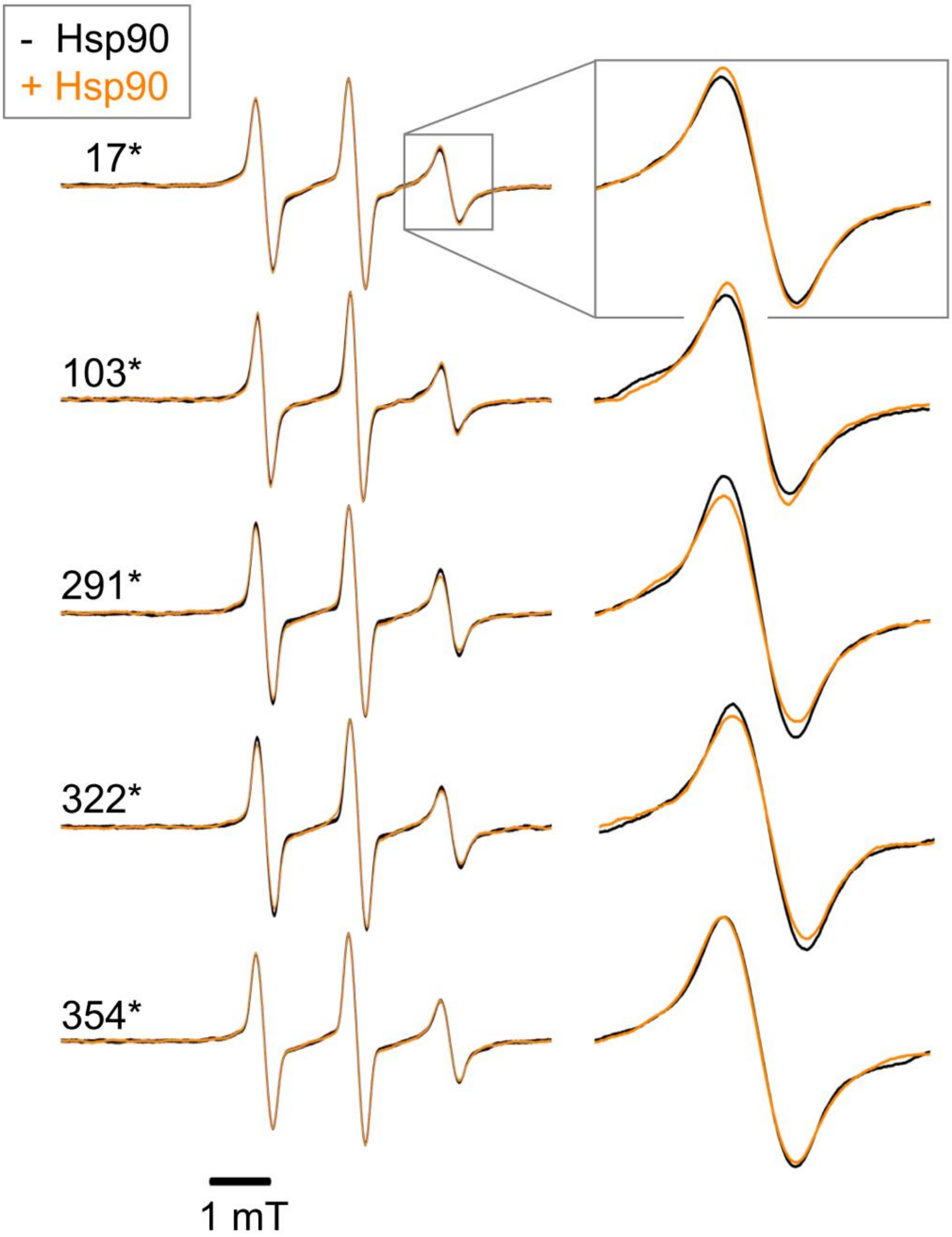

**Cw EPR data.** Cw EPR spectra of singly spin-labeled Tau in the absence (black) and presence of Hsp90 (orange). Close-up views of the high field spectral line are shown on the right for each Tau derivative. Spectral changes in the spectra of Tau upon addition of Hsp90 are very subtle, indicating that the Tau ensemble under observation is still highly dynamic also in the presence of the binding partner Hsp90. This is characteristic for a 'fuzzy' complex, which allows structural multiplicity and dynamic disorder while, nonetheless, the interaction between the binding partners is specific.

**Figure S9.**

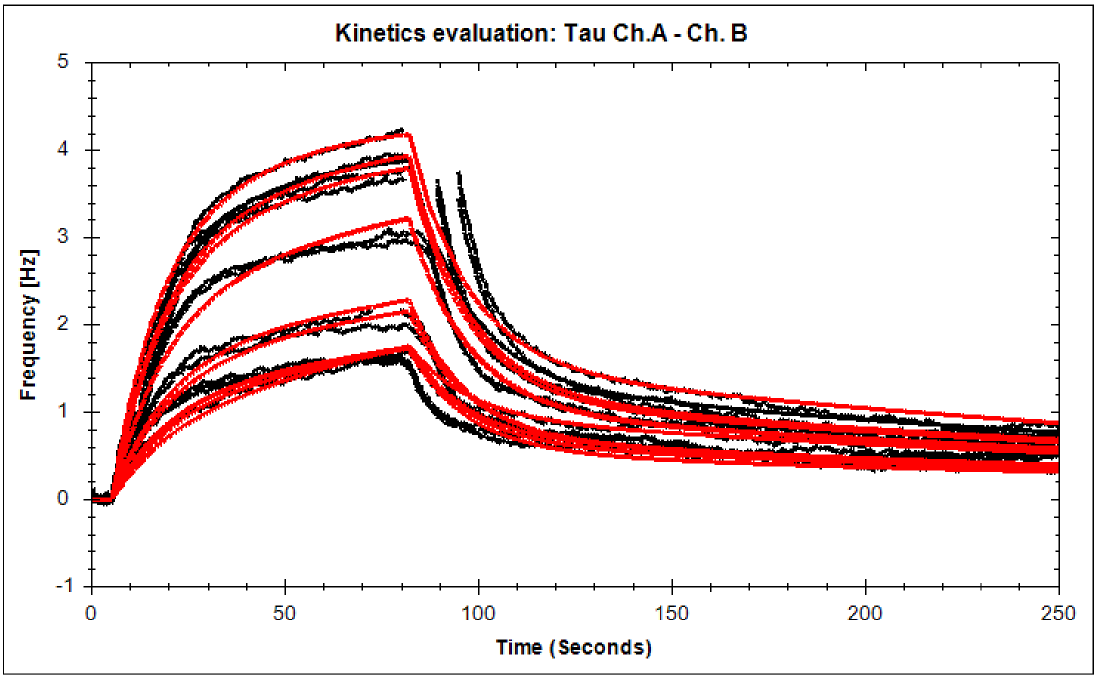

**Quartz crystal microbalance measurements.** QCM experiments were performed as described in materials and methods. The best fitting model (red lines) suggested 1:2 binding and thus two different modes of binding of Tau to Hsp90 as summarized in Table S2. 87% of Tau bound following the $k_{d1}$ characteristics in a transient way, and 13% of the binding followed the tight $k_{d2}$ characteristics. We explain the small contribution of the tight binding by the overall setup arrangement: Hsp90 was immobilized directly on the chip in small concentration, which might hinder the chaperone to maintain its dimer structure, leading to the exposure of some unusual binding surface. Thus, we reason that the $k_{d1}$ characteristics describe the Hsp90/Tau complex. The half lifetime of the Hsp90/Tau complex was estimated from the dissociation rate constant $k_{d1}$ as $\ln 2/k_{d1}$ and is approximately 10 s. This is characteristic of a transient protein-protein interaction.

### Tables S1 – S2

#### Table S1.

**Simulation of CD spectra.** Table S1 shows the respective percentages of secondary structural elements used in the simulation of CD spectra shown in Fig. S3.

| **Spectrum** | **α-helix** | **β-strand** | **β-turn** | **random coil** |
|---|---|---|---|---|
| Tau in sample buffer | 8.4 ± 5 % | 19.0 ± 5 % | 9.2 ± 5 % | 63.4 ± 5 % |
| Tau* in sample buffer | 8.4 ± 5 % | 19.0 ± 5 % | 9.2 ± 5 % | 63.4 ± 5 % |
| Tau in PBS buffer | 6.6 ± 5 % | 19.8 ± 5% | 6.6 ± 5 % | 67.0 ± 5 % |

#### Table S2.

**Kinetic analysis of QCM measurements.** Table S2 shows the kinetic analysis of QCM results as shown in Fig. S9.

| **Kinetic Analysis (1:2 Binding)** | | | |
|---|---|---|---|
| **$B_{max1}$** | **$k_{a1}$ (1/M·s)** | **$k_{d1}$ (1/s)** | **$K_{D1}$ (M)** |
| 87% | $2.46 \times 10^{+4}$ | $6.70 \times 10^{-2}$ | $2.72 \times 10^{-6}$ |
| **$B_{max2}$** | **$k_{a2}$ (1/M·s)** | **$k_{d2}$ (1/s)** | **$K_{D2}$ (M)** |
| 13% | $3.02 \times 10^{+4}$ | $3.49 \times 10^{-3}$ | $1.15 \times 10^{-7}$ |

### References

1. G. E. Karagöz *et al.*, Hsp90-Tau Complex Reveals Molecular Basis for Specificity in Chaperone Action. *Cell* **156**, 963-974 (2014).
2. S. Barghorn, P. Davies, E. Mandelkow, Tau Paired Helical Filaments from Alzheimer's Disease Brain and Assembled in Vitro Are Based on β-Structure in the Core Domain. *Biochemistry* **43**, 1694-1703 (2004).
3. H. H. de Jongh, E. Goormaghtigh, J. A. Killian, Analysis of circular dichroism spectra of oriented protein-lipid complexes: toward a general application. *Biochemistry* **33**, 14521-14528 (1994).
4. Y. Motozato, T. Nishihara, C. Hirayama, Y. Furuya, Y. Kosugi, Competitive inclusion of chloro-substituted acetic acids into β-cyclodextrin monitored by rotational correlation frequencies of 4-hydroxy-2,2,6,6-tetramethylpiperidinyl-1-oxy radical. *Can. J. Chem.* **60**, 1959-1961 (1982).
5. K. Krämer, *Physikalische Grundlagen der Maßeinheiten: Mit Einem Anhang über Fehlerrechnung*. (Springer-Verlag, 2013).
6. M. Pannier, S. Veit, A. Godt, G. Jeschke, H. W. Spiess, Dead-time free measurement of dipole–dipole interactions between electron spins. *J. Magn. Reson.* **213**, 316-325 (2000).
7. R. A. Stein, A. H. Beth, E. J. Hustedt, in *Methods Enzymol.,* Z. Q. Peter, W. Kurt, Eds. (Academic Press, 2015), vol. Volume 563, pp. 531-567.
8. S. Brandon, A. H. Beth, E. J. Hustedt, The global analysis of DEER data. *J. Magn. Reson.* **218**, 93-104 (2012).
9. D. Kurzbach *et al.*, Cooperative Unfolding of Compact Conformations of the Intrinsically Disordered Protein Osteopontin. *Biochemistry* **52**, 5167-5175 (2013).
10. D. Kurzbach *et al.*, Compensatory Adaptations of Structural Dynamics in an Intrinsically Disordered Protein Complex. *Angew. Chem. Int. Ed.* **53**, 3840-3843 (2014).
11. G. Jeschke *et al.*, DeerAnalysis2006—a comprehensive software package for analyzing pulsed ELDOR data. *Appl. Magn. Reson.* **30**, 473-498 (2006).
12. J. E. Kohn *et al.*, Random-coil behavior and the dimensions of chemically unfolded proteins. *Proc Natl Acad Sci U S A* **101**, 12491-12496 (2004).
13. A. M. Melo, J. Coraor, S. Elbaum-Garfinkle, A. Nath, E. Rhoades, A functional role for intrinsic disorder in the tau-tubulin complex. *Proc. Natl. Acad. Sci. U.S.A.* **113**, 14336-14341 (2016).
